# Circumvention of common labeling artifacts using secondary nanobodies

**DOI:** 10.1101/818351

**Authors:** Shama Sograte-Idrissi, Thomas Schlichthaerle, Carlos J. Duque-Afonso, Mihai Alevra, Sebastian Strauss, Tobias Moser, Ralf Jungmann, Silvio Rizzoli, Felipe Opazo

## Abstract

The most common procedure to reveal the location of specific (sub)cellular elements in biological samples is via immunostaining followed by optical imaging. This is typically performed with target-specific primary antibodies (1.Abs), which are revealed by fluorophore-conjugated secondary antibodies (2.Abs). However, at high resolution this methodology can induce a series of artifacts due to the large size of antibodies, their bivalency, and their polyclonality. Here we use STED and DNA-PAINT super-resolution microscopy or light sheet microscopy on cleared tissue to show how monovalent secondary reagents based on camelid single-domain antibodies (nanobodies; 2.Nbs) attenuate these artifacts. We demonstrate that monovalent 2.Nbs have four additional advantages: 1) they increase localization accuracy with respect to 2.Abs; 2) they allow direct pre-mixing with 1.Abs before staining, reducing experimental time, and enabling the use of multiple 1.Abs from the same species; 3) they penetrate thick tissues efficiently; and 4) they avoid the artificial clustering seen with 2.Abs both in live and in poorly fixed samples. Altogether, this suggests that 2.Nbs are a valuable alternative to 2.Abs, especially when super-resolution imaging or staining of thick tissue samples are involved.

## Main

Standard immunodetection approaches use typically a primary antibody (1.Ab) which binds the protein of interest (POI) and a secondary antibody (2.Ab) that binds to the 1.Ab and carries a detection element. In fluorescent microscopy techniques, the detection element is a fluorophore^1,2^ or a single strand of DNA. The latter is used in DNA Point Accumulation for Imaging in Nanoscale Topography (DNA-PAINT), a single molecule localization microscopy technique reaching <5 nm resolution by transiently binding of single stranded DNA bearing a fluorophore to their complementary strand on the target of interest^3,4^. The complex formed by the primary antibody and the secondary antibodies (1.Ab-2.Ab), is widely used because it is a cost effective and flexible approach since only 2.Abs need to be coupled to the detection element. However, the use of this complex carries some relevant limitations due to its size and due to the ability of 2.Abs to bind more than one epitope.

First, the 1.Ab-2.Ab can measure up to 30 nm, leading to a large distance between the targeted molecule and the detection element, causing the so called “linkage” or “displacement” error^5,6^. While this might not influence the results in some applications (e.g. epifluorescence, ELISA or FACS), it is of major relevance for super-resolution microscopy techniques where the localization precision can be as high as 1 nm^7,8^. The linkage error can be reduced by using directly labeled small affinity probes like camelid single domain antibodies (sdAbs) also known as nanobodies (Nbs)^5,9^, affibodies^10^, aptamers^11,12^ or affimers^13,14^, which all have sizes below 3 nm. Unfortunately, such small probes exist only for a handful of targets^15^, while conventional 1.Abs are easily available for a large number of POIs. An alternative to the standard 2.Abs was recently developed: monovalent recombinant secondary nanobodies (2.Nbs) which were reported to reduce the linkage error observed in dSTORM^16^.

Second, due to its steric hindrance the 1.Ab-2.Ab complex performs poorly in a crowded cellular environment or when the epitopes are abundant and densely arranged. In this respect, smaller probes such as aptamers or nanobodies are more efficient in the detection^11,17,18^. Moreover, sample penetration of full antibodies is a problem when staining thick biological samples such as tissues, biopsies or whole organisms^17,19^. For the optimal labelling of these thick samples, protocols have been established, but they are often laborious and require time-consuming incubations up to weeks^20^ or artifact-prone epitope retrieval protocols^21^. Smaller probes are expected to shorten the long incubation times.

Third, in multiplexed immunostaining, i.e. when multiple targets are stained in the same sample, one is constrained to the use of 1.Abs coming from different species. This is because the standard immunostaining needs to be done in a sequential manner: first 1.Abs are incubated on the sample, washed off and then 2.Abs are incubated. Therefore, 1.Abs should be raised in different species and 2.Abs should recognize one species specifically, limiting the choice of antibodies for the different targets. It has been shown that by pre-mixing 1.Abs with 2.Nbs in a tube prior staining, one could circumvent this species limitation and use on a sample 1.Abs raised in the same species^16^.

Finally, conventional antibodies used for immunodetections are bivalent binders, *i.e.* each antibody can recognize two POIs/epitopes simultaneously. Moreover, 2.Abs are commonly a mixture of binders which can bind to different epitopes of the same 1.Ab (polyclonal). The combined bivalency and polyclonality of antibodies used in standard immunodetection procedures have been shown to induce clustering of the POI and their interactors seriously affecting some conclusions based on such method^22,23^. Using a monovalent secondary probe these clustering effects might be reduced.

In this work we tested and thoroughly validated the use of 2.Nbs for several microscopy applications. We first confirmed that the usage of 2.Nbs decreases linkage error in STED microscopy and DNA-PAINT. We then exploited the ability of those probes to allow the simultaneous use of several 1.Abs from the same specie by using them in 3D Exchange-PAINT. This technique enables to image a virtually infinite number of targets in high resolution in the same sample^24,25^. We show that 2.Nbs are the optimal tools to complement the multiplexing capability of this super-resolution technique. Additionally, we observed that the pre-mixing of 1.Ab-2.Nb can save time in staining thick biological samples imaged under light sheet microscopy, ensuring also a better sample penetration and homogenous staining. Finally, we systematically compared the probe-induced clustering of the target protein either using directly-labeled monovalent probes, like affibodies and single Fab’ fragments, and conventional 1.Abs detected by polyclonal and bivalent 2.Abs or by 2.Nbs. We observed that 2.Nbs drastically reduced the probe-induced clustering of the target in both live and fixed sample. This makes 2.Nbs a real alternative to conventional 2.Abs by minimizing experimental time, expanding the multiplexing ability of immunostainings, improving the tagging precision and signal linearity, and finally avoiding the probe-induced clustering artifacts.

## Results

### Secondary nanobodies provide higher staining accuracy than secondary antibodies

First, we investigated the accuracy of 2.Abs or 2.Nbs in revealing their target 1.Ab. To do so, we imaged in a Two color STED microscopy setup COS-7 cells stained with a monoclonal 1.Ab anti-tubulin directly conjugated to AbberiorStar635P and further recognized by either a polyclonal 2.Ab or a monovalent 2.Nb both carrying AbberiorStar580. An autocorrelation analysis was performed on these images to evaluate the staining accuracy of the secondary probes. Initially, the autocorrelation of the 1.Ab images provided an idea of the distribution or density of the 1.Ab on microtubule filaments. The autocorrelation curve obtained from the signal of the 2.Nb followed the trend of the autocorrelation obtained for the 1.Ab, which proposes that the 2.Nb signal accurately follows the fluorescent signal from the 1.Ab. In contrast, when performing the same analysis on the staining performed with polyclonal 2.Abs, the correlation curve was shifted to the right. This suggests that the 2.Ab inaccurately reveals the location of the 1.Ab (Fig 1A). To confirm this, we decided to look at peroxisomes within primary hippocampal neurons. We compared the diameter of these small organelles when imaged with STED microscopy after using a 1.Ab revealed by a 2.Ab or 2.Nb. We observed a clear and significant shift to smaller diameters of these organelles after comparing 3020 peroxisomes stained with 1.Ab-2.Ab and 3109 peroxisomes stained with 1.Ab-2.Nb (Fig 1B).

**Figure 1.**
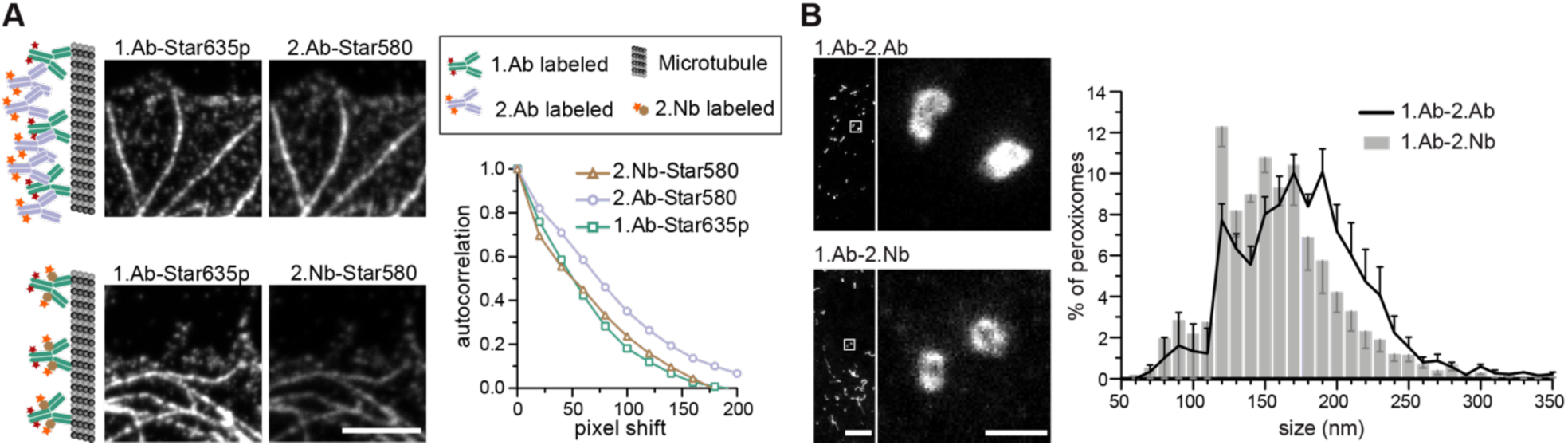
2.Nbs minimize the linkage error caused by 2.Abs and increase detection accuracy. (**A**) Two color STED imaging of microtubules stained with 1.Ab directly labeled with AbberiorStar635p dye and secondary reagents (either 2.Ab or 2.Nb) labeled with AbberiorStar580. Example images and schematic representation of the experimental procedure. Scale bar 2.5 µm. Autocorrelation analysis on signal obtained from either the 1.Ab or the secondary probe microtubules. N=51 line profiles for 2.Nb, N= 70 for 2.Ab and N=121 for 1.Ab. One-way ANOVA p=1.061×10^−6^ F=14.58 followed by post hoc Bonferroni tests indicates that the 2.Ab is different with p<0.01 from the 1.Ab and 2.Nb which themselves are indistinguishable. (**B**) Neuronal peroxisomes stained with 1.Ab-2.Ab or 1.Ab-2.Nb. Exemplary STED images. Scale bars 10 µm (overview) and 100 nm (zoom). Size distribution analysis of peroxisome. N=3109 peroxisomes were analyzed when stained with 2.Nb and N= 3020 stained with 2.Ab. Error bars correspond to the standard error of the mean from 4 independent experiments. Paired t-test shows that the apparent size of peroxisomes stained with 2.Nb is on average smaller with p<0.01 compared to the one stained with 2.Ab.

To more precisely evaluate if the 2.Nbs decrease the linkage error, we required higher resolution capability. For this purpose, we used DNA-PAINT that has achieved resolution below 10 nm^4^. DNA-PAINT uses affinity reagents attached to short DNA oligonucleotides, so we first coupled the 2.Nbs to a single stranded DNA oligo (termed docking strand) as described previously^26^. We performed an assay which has been used as gold standards in the field to assess linkage error^27^. We stained the microtubule network of a fibroblast cell line with a monoclonal 1.Ab against alpha-tubulin detected with a 2.Nb coupled to a docking strand (Fig. 2A). We measured an apparent diameter of ∼30 nm for a microtubule filament in a 2D projection (Fig. 2F) which is smaller when compared to microtubules stained using 1.Ab and 2.Ab (Fig. 2C) revealing a filament width of ∼50 nm.

**Figure 2:**
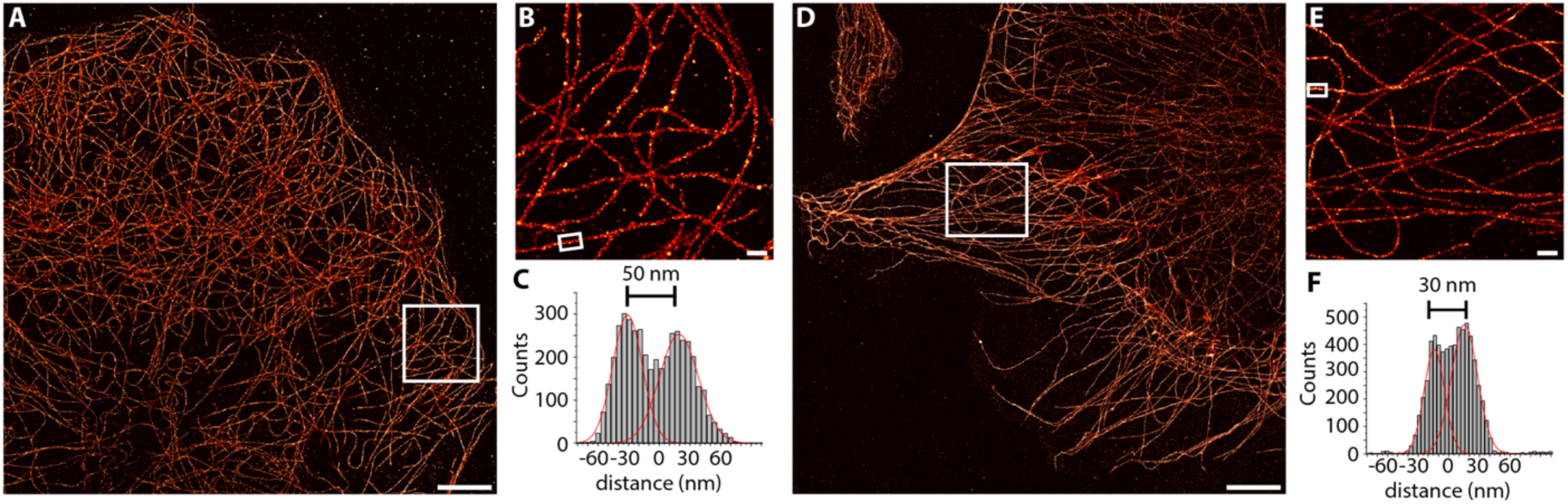
DNA-PAINT imaging with 2.Nbs confirms that 2.Nbs minimize the linkage error caused by 2.Abs. (**A**) Overview DNA-PAINT image of COS-7 cells stained with 1.Ab targeting alpha tubulin and 2.Ab coupled to a DNA-PAINT docking strand. (**B**) Zoom-in image of the region highlighted in A. (**C**) Cross-sectional histogram analysis of the highlighted region in B results in a microtubule filament diameter of ∼50 nm. (**D**) Overview DNA-PAINT image of alpha tubulin stained with 1.Ab and 2.Nb (**E**) Zoom-in image of the region highlighted in D. (**F**) Cross-sectional histogram analysis of the highlighted region in E results in a smaller microtubule filament diameter of ∼30 nm. Scale bars: 5 μm (A, D), 500 nm (B, E).

### Bypassing the primary antibody animal-species limitations

In standard immunoassays, normally the 1.Abs are first incubated with the sample followed by washes to eliminate the non-bound excess of 1.Abs. Only at this point 2.Abs are incubated for a period of time with the sample followed by washes to eliminate the non-bound excess of 2.Abs before imaging the specimen. Pre-mixing the 1.Ab with the 2.Ab prior to incubating them with the sample would shorten protocols and save considerable amount of time and costs (*e.g*. in clinical pathology laboratories). However, this is currently not possible due to the polyclonality and the bivalency of 2.Abs that result in agglutination (aggregation or clustering) of the 1.Abs-2.Abs complex and thus in a failure to stain the intended target in the sample (Supp. Fig. 1, left panel). If the secondary probe binds to the 1.Ab in a monovalent fashion, pre-mixing primary and secondary probes is possible. The pre-mixing of 2.Nbs with a mouse monoclonal 1.Ab against alpha tubulin in a tube for 15 minutes resulted in properly stained filaments (Supp. Fig. 1, right panel) or single bands detected in a fluorescent Western blot assay (Supp. Fig. 2A). Bypassing this pre-mixing limitation with monovalent secondary probes open a new possibility previously not possible in immunoassays: using several 1.Abs raised from the same species. Conventional immunoassays that allowed the detection of two or more POIs require that each 1.Ab has to come from a different animal species (e.g. mouse, rabbit and chicken for the detection of 3 POIs on the same specimen). This strict requisite is necessary to ensure the indirect detection of the POIs with species-specific 2.Abs. This restriction provides a great limitation for the choice of 1.Ab and reduces the multiplexing capability of any immunoassay. Here we chose three different monoclonal 1.Abs raised in mouse directed against alpha-tubulin, GM130 (Golgi), and FXFG repeats in nucleoporins (nuclear pore complex; NPC). Each was pre-mixed with 2.Nbs anti mouse carrying each a different fluorophore (Fig. 3B). The COS-7 cells imaged under scanning confocal microscopy clearly displayed the three stained structures (microtubules, Golgi and NPC) with minimal background and negligible cross-talk between the channels. Similarly, the pre-mixing capability of these 2.Nbs using the same species of 1.Abs for multiple targets could also be applied for the detection of 2 different POIs in fluorescent Western blots assays (Supp Fig. 2B).

**Figure 3.**
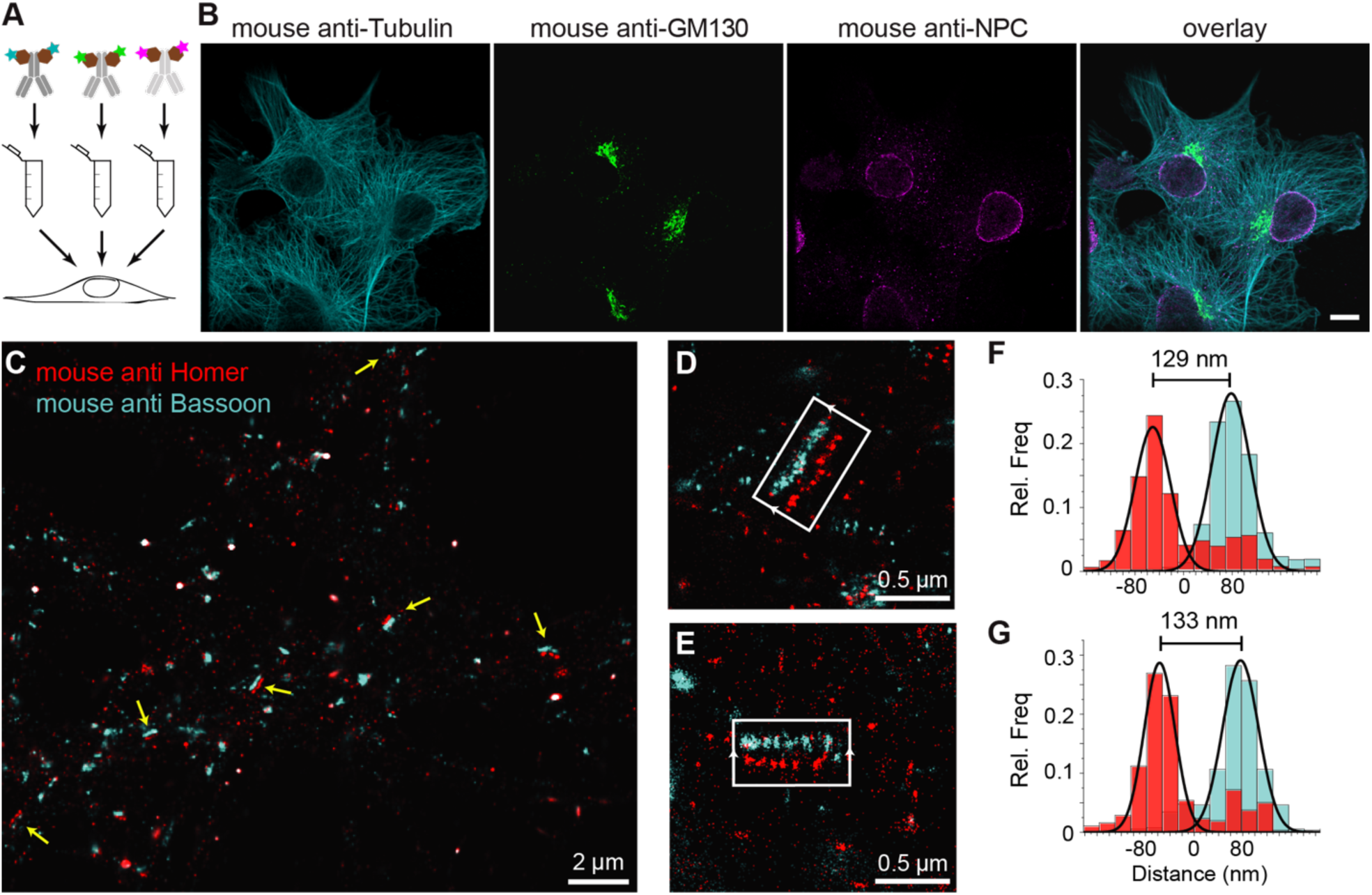
Pre-mixing 1.Abs with 2.Nbs allows to use same animal-species 1,Ab for several target proteins on the same sample. (**A**) Scheme of pre-mixing: different 1.Abs were pre-mixed with 2.Nbs each carrying different fluorophores and subsequently incubated on cells. (**B**) Example of confocal images performed on a sample stained with the pre-mixing methods. Cyan: mouse anti-Tubulin 1.Ab pre-mixed with 2.Nb-CF633. Green: mouse anti GM130 1.Ab pre-mixed with 2.Nb-Alexa488. Magenta: mouse anti NPC 1.Ab pre-mixed with 2.Nb-Alexa 546. Scale bar represents 10 µm. (**C**) 3D Exchange-PAINT overview image of primary rat hippocampal neurons. Yellow arrows indicate evident mature synapses where the pre-synaptic active zone (mouse 1.Ab anti Bassoon) and post synaptic density (mouse 1.Ab anti Homer) are in front of each other. (**D,E**) higher magnification of 2 selected synapses where a clear synaptic cleft is recognized. (**F, G**) histogram analysis of the selected synapses displaying the length of the synaptic cleft.

The pre-mixing feature seems to be ideal for a technique that allows the detection of multiple targets (multiplexing). Therefore, we turned once again to DNA-PAINT, this time we used an extension termed Exchange-PAINT which can, in theory, image an unlimited number of POIs on the same sample with a few nanometer precision^24,26^. We stained primary hippocampal neurons with two mouse monoclonal 1.Abs and each was pre-mixed with 2.Nbs conjugated to DNA docking strands with orthogonal sequences. We performed 3D Exchange-PAINT on synapses stained against Bassoon, a protein highly enriched at the presynaptic active zone^28^, and the scaffold protein Homer that is concentrated at the postsynaptic density^29^ (Fig.3C). Notably, we obtained a super-resolved view of a single neuronal synapse in 3D using two 1.Abs from the same species. We estimated a distance of 130 nm (Fig.3D-G) between bassoon (presynaptic) and homer (postsynaptic) reproducing findings made with other advanced microscopy techniques such as dSTORM^30^ and X10 expansion microscopy^31^.

### Secondary nanobodies enhance sample penetration in shorter incubation time

We used the time advantage of pre-mixing 1.Ab with the 2.Nb in a complex thick sample which requires long incubation with the probe to ensure proper sample staining. We used cochleae extracted from three weeks old mice and stained parvalbumin-*α*, a calcium buffering protein present in inner hair cells and type I spiral ganglion neurons (Fig. 4). We compared how long the 1.Ab-2.Nb and 1.Ab-2.Ab needed to be incubated to ensure a homogenous staining throughout the sample. In order to image the entire volume, we used light sheet microscopy after decalcification and clearing. Two cochleae obtained from the same animal were stained either with 1.Ab and sequential 2.Ab or with 1.Ab pre-mixed with 2.Nb for comparable amount of times. The cochleae stained with 1.Ab-2.Ab for 6 days (3 days 1.Ab, 3 days 2.Ab) showed insufficient penetration of the staining, with signals accumulated in the outer bone surface and in the edges exposed to the solution (Fig. 4A). The cochlea stained with the same antibody for 14 days (7 days 1.Ab, 7 days 2.Ab) showed a better staining performance, revealing hair cells and neurons. However, the ganglion displayed a staining gradient with stronger signals on the edges, indicating insufficient detection of target molecules (Fig. 4A). On the other hand, the cochleae stained with pre-mixed 1.Ab-2.Nb for 6 and 14 days revealed a homogenous staining of neurons. No apparent difference between the two incubation times was observed (Fig. 4A). An incubation lasting less than 6 days might be even sufficient. A custom written analysis quantifying the signal intensity throughout the ganglion of the cochleae (Fig. 4C, Supp. Figs 3 and 4) showed how the signal coming from cochleae stained with 1.Ab-2.Nb displayed a plateau phase meaning homogenous staining, while the ones stained with 1.Ab-2.Ab displayed a peak showing a gradual staining from the distal to the central portion of the ganglion.

**Figure 4:**
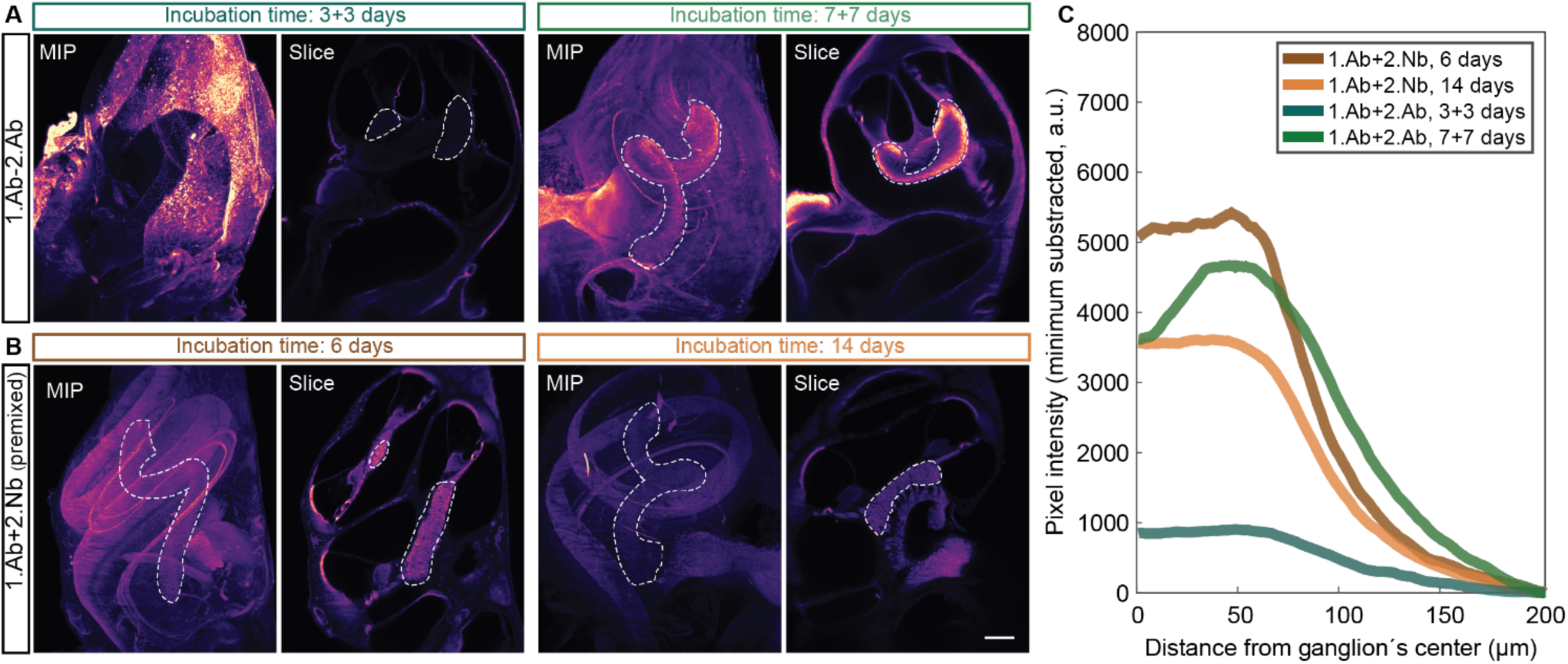
Pre-mixing decreases the incubation time necessary to obtain homogenous staining of the cochlea. Mice intact cochlea were stained with a parvalbumin-*α* antibody, either pre-mixed with 2.Nb (**A**) or sequentially incubated with 2.Ab (**B**) for the time indicated. In each panel the maximal intensity z-projection (MIP) and an exemplary light sheet microscopy slice of an intact cochleae (Slice) are depicted. Scale bar: 200 µm. Ganglion outlined by dotted line (**C**) Mean pixel line profile from radii crossing the ganglion distributed along the centerline of the ganglion. See Supp. Fig. 3 for schematic analysis explanation and Supp. Fig. 4 for raw data. N=2 cochlea per condition. Note: the plateau profile depicted by the samples stained with 2.Nb, as opposed to the relatively pronounced peak profile in the samples stained with 2.Ab for 7 days or to the flat profile in the samples stained for 3+3 days.

### Secondary nanobodies reduce probe-induced clusters of target proteins on living cells

To test if the 2.Nbs reduce probe-induced clustering of target molecules, we decided to analyze the surface distribution of IgM containing B cell receptors (IgM-BCRs) on a human B cell line (Ramos cells). This cellular model allows simple visual inspection and numerical analysis because the POIs are evenly distributed in the cellular surface of these resting B cells^32^. Cells were stained and chemically fixed with aldehydes to be imaged under stimulation emission depletion (STED) microscopy. Initially, the surface IgM-BCRs on living Ramos cells were stained using monovalent probes: a monoclonal affibody^32^ or a polyclonal single Fab’ fragment (polyFab’) (Fig. 5). In this case a smooth continuous plasma membrane signal from the surface IgM-BCRs was observed at the optically sliced equator of the cells. However, when cells were stained using a 1.Ab-2.Ab, a sparse clustered signal was clearly identified (Fig. 5).

**Figure 5.**
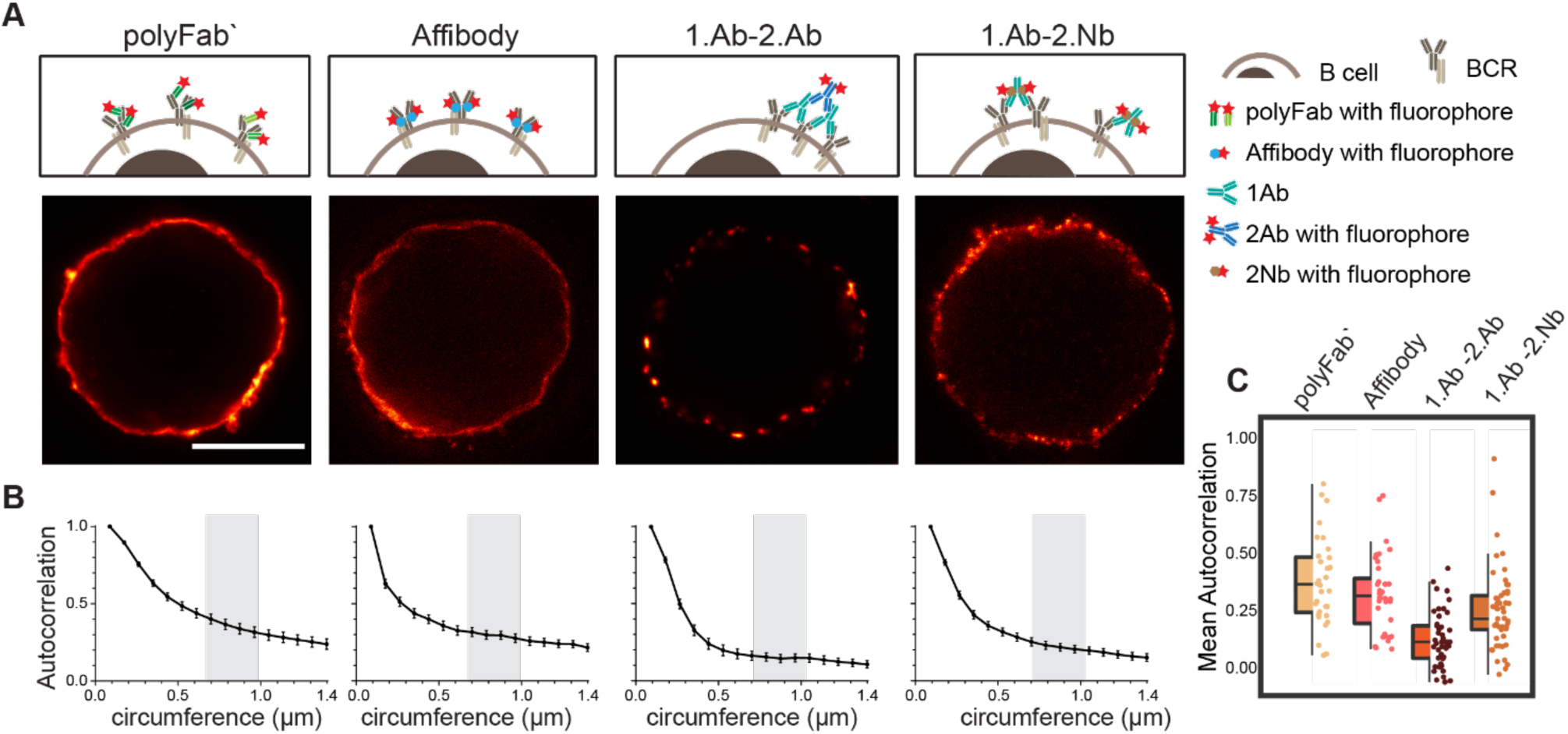
Live staining of IgM-BCRs on Ramos cells shows different pattern according to the probe used. (**A**) STED images of a B cell stained with fluorescent polyclonal single Fab’ fragment (polyFab’); affibody; primary antibody revealed by a fluorescent secondary antibody (1.Ab-2.Ab); primary antibody revealed by a secondary nanobody (1.Ab-2.Nb). All fluorescent probes were conjugated to AbberiorStar635P fluorophores. Scale bars = 5 µm. (**B**) Autocorrelation analysis along the circumference of cells. We analyzed three independent experiments with N≥ 10 cells for each conditions (**C**) Box-dot plots show the average autocorrelation from 0.7 µm to 1.0 µm circumference (grey zones shown in the graphs in **B**). Boxes show the interquartile range (IQR). Lines signify medians, and whiskers extend to 1.5 times the IQR. Lower box represents higher clustering. The autocorrelation observed in 1.Ab-2.Ab differs from the monovalent probes (polyFab’ and affibody) by P ≤ 0.0001 and from 1.Ab-2.Nb by P ≤ 0.001. P values were calculated with one-way ANOVA followed by Tukey Multiple Comparison Test. See Supp. Table 5 for full statistics.

Finally, we tested if the 2.Nbs elicit a similar clustering effect observed using 1.Ab-2.Ab detection system. Interestingly, a considerably milder effect was observed when using the same 1.Ab detected by a 2.Nbs, partially rescuing the pattern observed with the monovalent affibody or the polyFab’ that bind directly to the IgM-BCRs (Fig. 5). This result suggests that although the bivalency of the monoclonal 1.Ab still deviates slightly from the signal distribution obtained with fluorescent monovalent primary probes, the major cluster-inducing element seems to be contributed by the polyclonality of the 2.Abs. A Pearson‘s autocorrelation analysis^33^ was used to quantify the probe-induced clustering. The custom-written analysis consists of collecting the STED image intensity along the membrane and correlating it to itself for different rotation angles. We then plotted the autocorrelation curves, which start with a perfect correlation (*r* = 1) at zero rotation and decrease at higher rotations (Fig. 5B, with rotation angle converted to corresponding membrane distance). The major empirical effect between the different conditions was observed at membrane distances between 0.7 to 1 µm. Therefore, the correlations measured throughout this interval were then averaged with the value obtained from each cell corresponding to a spot on the scatter plot (Fig. 5C). With this method we determined an average autocorrelation of 0.34±0.19 (mean±SD) for cells stained with the monovalent polyFab’ and 0.29±0.16 for cells stained with the monovalent affibody, while it was only 0.07±0.12 for the cells stained with monoclonal 1.Ab and polyclonal 2.Ab. These analysis quantify the probe-induced clustering of IgM-BCRs observed by the 1.Ab-2.Ab condition. Interestingly, this effect was not only evident using super-resolution microscopy, but it was also observed in diffraction limited scanning confocal microscopy images (Supp. Fig. 5). As expected by the more continuous pattern observed, the average autocorrelation of cells stained with 1.Ab-2.Nb was 0.21±0.17 indicating a significant decrease (rescue) of the probe-induced clustering artifact caused by the polyclonal 2.Ab (Fig. 5).

### Probe-induced clusters of target proteins in aldehyde-fixed cells

It has been noticed that conventional fixations times with 4% paraformaldehyde (PFA) does not necessarily prevent protein movement^34^. Also other variables like blocking reagents and temperature need to be taken into consideration and tested case-by-case depending on the imaged target^35^. A more efficient fixative such as glutaraldehyde (GLU) could be used, but it generates unwanted autofluorescence and only few affinity molecules find their target epitopes after GLU crosslinking. A recently described di-aldehyde alternative that seems to alleviate some of these problems caused by PFA and GLU is glyoxal^36^, although its implementation is very recent and the vast majority of researchers still use PFA-fixation for conventional immunofluorescence. Therefore, we tested and compared the probe-induced clustering after exposing the Ramos cells for 10 and 30 minutes with 4% PFA or 30 minutes with a combination of 4% PFA and 0.1% GLU (Fig. 6 and Supp Fig. 6). We compared these fixative condition and live staining using the classical 1.Ab-2.Ab complex or the 1.Ab-2.Nb imaged under STED microscopy. Our observations suggest that applying 4% PFA for 10 minutes is not enough to avoid the artifactual clustering formation induced by 1.Ab-2.Ab (autocorrelation of 0.14±0.11 not significantly different from the live staining condition 0.07±0.12). However, 4% PFA for 30 minutes seems to be sufficient to rescue to a great degree the clustering artifact caused by the 2.Ab (0.23±0.18 different from the live staining condition with p value P ≤ 0.001; see also Fig.2A). Using the 2.Nbs had no significant change between live, 10 or 30 minutes of fixation with 4% PFA (0.21±0.17, 0.20±0.17 and 0.21±0.18 respectively; Fig. 6B). As expected, similar non-clustering effects are observed for a primary monovalent probe like the polyFab’ directed against human IgM-BCRs (Supp. Fig. 6). In addition, when observing the staining pattern created by the combination of 4% PFA and 0.1% GLU for 30 minutes, the stained rim of the cells is not a thin layer as observed by PFA fixation, but it displays a texture-like surface. From studies in electron microscopy, it is expected that GLU fixation results in a better ultrastructure preservation and in this case. Due to the uneven texture-like surface when fixing with PFA and GLU, and therefore a reduced homogeneity at the investigated spatial scale, the Pearson‘s correlation analysis has the tendency to paradoxically display a slightly lower correlation (Fig. 6-boxplot).

**Figure 6:**
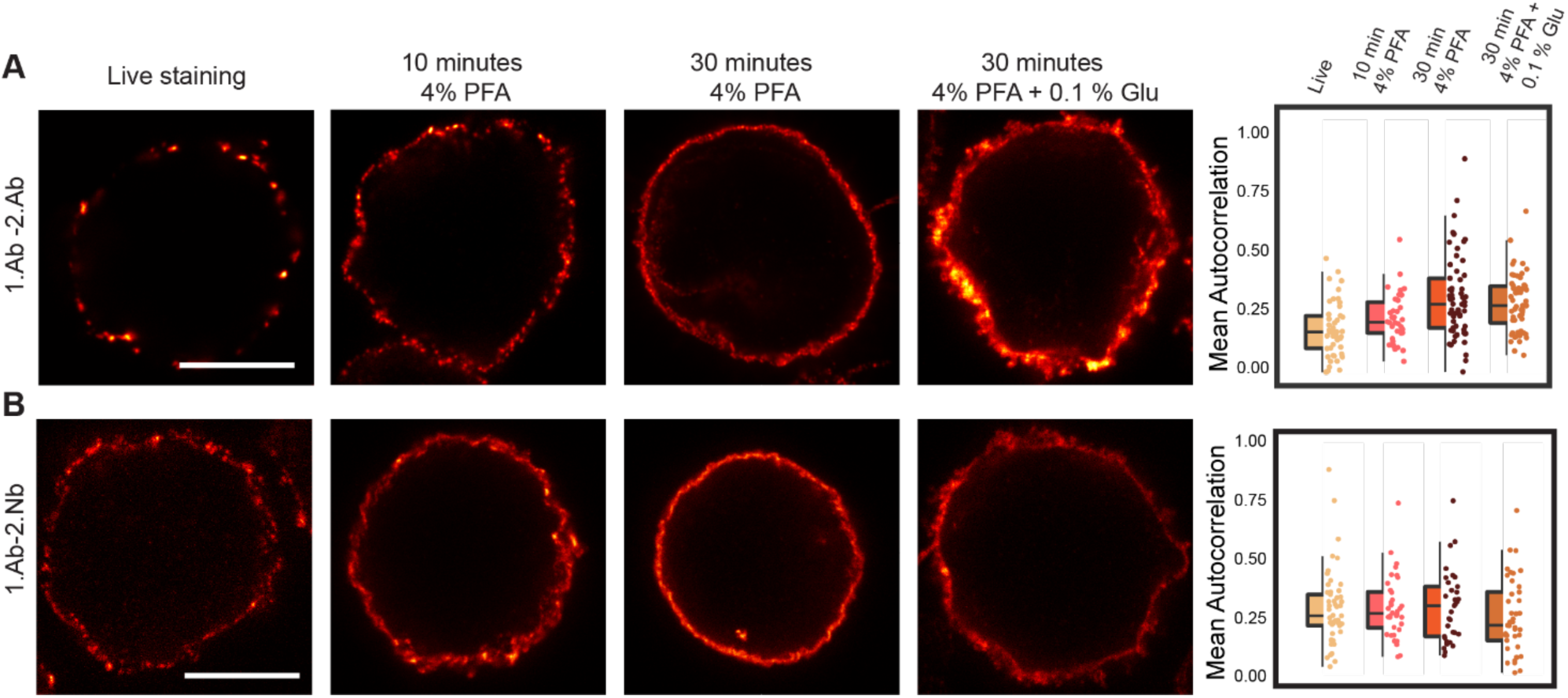
Probe-induced clustering on aldehyde fixed B cells. (**A**) and (**B**) STED images showing the effect of fixation on clustering induced by 1.Ab-2.Ab or 1.Ab-2.Nb. Scale bars represent 5 µm. Box-dot plots on the right represent the autocorrelation averages and are obtained as in **Fig.4C**. Autocorrelation curves are detailed in Supp. Fig. 7 and statistics in Supp. Table 5

## Discussion

In this study we have shown how 2.Nbs can be used to overcome some limitations of and artifacts caused by the use of conventional polyclonal 2.Abs. Direct labeling of the 1.Ab with a reporter avoids the use of a secondary probe. However, this approach is expensive since it requires a large amount of starting material and most of the labelling strategies can result in antibodies not binding their target. Major efforts have been taken to increase the reproducibility in science and it has been pointed out that an important contribution in biological applications arise from poorly characterized antibodies (especially polyclonal serums, which can be highly heterogeneous)^37^. Additionally, using recombinantly produced 2.Nbs not only increases the reproducibility of biomedical experiments, but importantly it eliminates the ethically controversial use of animals for 2.Abs (donkey, goat, sheep, etc). Moreover, their recombinant production allows a high level of flexibility to modify them in a controlled manner, for example with site-specific and quantitative conjugation of flurophores^9^ or docking strands for DNA-PAINT^26^.

### Smaller size of the secondary probe decreases linkage error and increases staining accuracy

We have shown that the use of 2.Nbs offers extra advantages due to their small size. It has been already suggested that nanobodies binding directly the POI were able to minimize the delocalization of the fluorophores with respect to the POIs^5,9,38^. Here, we can show using STED on peroxisomes of primary neurons and DNA-PAINT analysis of microtubules network on fibroblasts (Fig. 1B, Fig. 2) that the linkage error provided with 2.Abs is also reduced by replacing them with 2.Nbs. Additionally, we found that 2.Abs represent poorly the accurate location of the 1.Abs (Fig. 1A). We show that the 1.Ab directly labeled with a fluorophore decorates microtubules with a certain periodicity that can be followed when revealed with a 2.Nb but not with a conventional 2.Ab, suggesting that the 2.Ab blurs the localization of the 1.Ab. This inaccuracy of the polyclonal 2.Abs has a major consequence in one of the main application of fluorescence microscopy: protein co-localization studies. Fluorescently labeled 2.Abs can also mis-represent the quantities of 1.Abs and in turn wrongly estimate the levels of the POIs. For instance, the same amount of target proteins in two different spots might be erroneously under- or over-represented by the number of 2.Abs decorating the 1.Abs and by the random localization/number of the fluorophores on these 2.Abs. This can be better controlled using monovalent binders carrying a controlled number of fluorophores, making signal intensities in microscopy more linear with respect to amount of target protein.

### Pre-mixing overcomes the species limitation for multiplexing microscopy

Mixing the primary and the secondary reagents prior to incubating them with the sample (pre-mixing) is a desired feature as it saves experimental time. Unfortunately, it cannot be performed with conventional bivalent polyclonal 2.Abs (Supp. Fig. 1), however, pre-mixing works when using monovalent binders against 1.Abs^16^. More importantly, pre-mixing eliminates the animal-species limitation of the primaries when detecting two or more POIs. We first showed using a conventional three channel/colors scanning confocal microscopy that it is possible to use three mouse 1.Abs in the same sample. Nevertheless, pre-mixing needs to be tested and well validated for every particular set of 1.Abs, since the 2.Nbs are not covalently bound to the 1.Abs. Therefore, there is a risk of 2.Nbs unbinding from the intended 1.Ab and binding to a different 1.Abs present in the sample. This would cause a cross-contamination of signals. For that reason, we have decided to perform a short post fixation between each color as an strategy to ensure the immobilization of the 2.Nbs, but to maintain the animal-species freedom in demanding multiplexing super-resolution imaging. Here we showcased the proof-of principle of pre-mixing immunostaining for 3D Exchange-PAINT super-resolution microscopy. We determined the distance between the pre- and post-synapses (synaptic cleft) with a good accuracy using two 1.Abs raised in mice. Imaging these well-studied structures helped to demonstrate that negligible signal contamination was observed using our pre-mixing approach. Exchange-PAINT not only provides impressive resolution and allows 3D imaging, but it also eliminates the limit on the number of POIs that can be imaged in the same sample. This makes the combination of pre-mixing using 2.Nbs with Exchange-PAINT a very powerful multiplexing technique with 3D and exquisite spatial resolution capabilities.

### Pre-mixing shortens experimental time and allows a better penetration of probes in thick tissue

Immunostaining protocols of complex thick tissue samples typically require days to weeks^20,39^, especially since the 1.Abs and 2.Abs have to be added sequentially. Pre-mixing methodology reduced the time of all immunostainings, but it is clearly a time-saver when used in long staining protocols. Here we used cleared mouse cochlea imaged with light sheet microscopy to compare the staining pattern if using sequential 1.Ab and 2.Ab or pre-mixed 1.Abs with 2.Nbs. Our observation suggest that pre-mixing shorten the conventional protocol by at least half the time (*i.e.* 6 days of staining; Fig. 4). We did not test shorter times for pre-mixing, but the fact that no clear difference in intensity or signal distribution between pre-mixed stainings for 6 or 14 days were observed, suggests that optimal incubation time might be even shorter. Additionally, pre-mixing using 2.Nbs provided a specific and homogenous staining regardless of the depth of the target parvalbumin-*α* positive cells (Fig. 4). This was not the case with conventional antibodies, where the staining after 6 days was primarily on the surface and although at 14 days some penetration occurred, it displayed a clear disproportional staining (not homogenous), suggesting that more parvalbumin-*α* is at the surface that inside the spiral ganglion. Since levels of parvalbumin-*α* are comparable among all the different subtypes of type I spiral ganglion neurons^40^ and type I spiral ganglion compose 96% of the neuronal population of the ganglion^41^, this is not expected to be true, but in turn, to be an artifact of penetration or steric hindrance of the antibodies.

### Antigen clustering on cells rescued by the use of secondary nanobodies

In this study we have shown how conventional polyclonal 2.Abs can induce the clustering of 1.Abs and thus the clustering of target proteins (the antigen). Our study also provides an alternative to rescue this probe-generated artifact. This can be achieved by either specific monovalent affinity tools binding directly the POIs (like nanobodies, affibodies, single Fab’ fragments, etc) or by replacing the polyclonal bivalent 2.Ab with monoclonal and monovalent 2.Nbs (Fig. 5, Fig. 6). Staining living cells show clearly how the distribution of BCRs went from sparsely and homogenously distributed when using a monovalent primary probe such as an affibody or a single Fab’ fragment, to a clustered pattern when using 1.Ab-2.Ab. This can be expected since the bivalency and the polyclonality of the two combined probe can force clustering of 1.Abs and thus of target proteins. The combination of 1.Ab-2Nb on the other hand seemed to rescue this artifact to a good degree and showed significantly less probe-induced clusters (Fig. 6), suggesting that the monoclonal bivalent 1.Ab has some minor effect on the clustering compared to the major clustering observed when using the polyclonal 2.Ab.

Importantly, this probe-induced clustering artifact could also be observed when short aldehyde fixation was performed. Sample fixation with 4% paraformaldehyde for 10 minutes, is a widespread practice in biology laboratories, but seems to be insufficient to fully immobilize cellular elements. Although it was previously suggested that short aldehyde fixation is not sufficient to stop molecular movement^42^, we observed this time that it is actually also not sufficient to prevent the 2.Abs to induce clustering of the POIs (Fig. 6). The artificial aggregation of POIs even after chemical fixation can lead to several misleading conclusions when studying for example co-localization of two or more POIs, poly-molecular arrangements or if molecular mechanisms are interpreted after imaging analysis. Increasing the time in which the aldehyde fixation is applied to 30 minutes, reduced the 1.Ab-2.Ab induced clustering of BCRs significantly. The type of fixative and its application time seemed not to influence the location detection of the BCR when stained with a monovalent primary probe such as Fab’ fragment (Supp. Fig. 6) or with 1.Ab-2.Nb (Fig. 6). Therefore we conclude that poor fixation in combination with bivalent and polyclonal affinity tools like conventional secondary antibodies might result in an artificial distribution of the proteins under study.

Small, monovalent, monoclonal probes specific to their target are clearly the ideal probe to reveal the POIs. Unfortunately, their availability is limited to a handful of targets. On the other hand, a large amount of well validated monoclonal antibodies is available. Our data suggest that if 1.Abs detected with recombinant 2.Nbs (and probably any other small monovalent binder such as Protein A and Protein G^43^), can minimize the artifactual clustering of target, increase the localization accuracy in super-resolution microscopy, lower steric hindrance to detecting more target molecules, increasing the sample penetration, multiplexing staining capabilities and finally increase the reproducibility with no needs of animals can be envisioned.

## Material and Methods

### Cell culture

Cells were cultured at 37°C, 5% CO_2_ in a humified incubator. The human Burkitt lymphoma B cell lines DG75 and COS-7 fibroblast were obtained from the Leibniz Institute DSMZ—German Collection of Microorganisms and Cell Culture (DSMZ Braunschweig, Germany). For maintenance, cell lines were kept on petri dishes. For experiments cells were plated on poly-L-lysine (PLL)-coated coverslips. DG75 cells were splitted every 3 days using fresh complete medium (RPMI medium supplemented with 10% Fetal Bovine Serum (FBS), 4 mM L-glutamine and 100 U/ml penicillin and streptomycin). COS-7 fibroblast cells were cultured in complete Dulbecco’s MEM with the addition of 10% FBS, 4 mM L-glutamine, 0.6% penicillin and streptomycin. A549 cells (ATCC, Cat. No. CRL-1651) were maintained in DMEM (Thermo Fisher Scientific, Cat. No. 10566016), supplemented with 10% Fetal Bovine Serum (Thermo Fisher Scientific, Cat. No. 10500-064) and 1% Penicilin/Streptomycin (Thermo Fisher Scientific, Cat. No. 15140-122). Rat primary hippocampal neuron cultures were prepared as described before by Opazo *et al*.^33^ In brief, the brains of P1-2 were extracted and placed in cold HBSS (ThermoFisher, Waltham, Massachusetts, USA). The hippocampi were extracted and placed in a solution containing 10 mL DMEM (Thermo Fisher), 1.6 mM cystein, 1 mM CaCl_2_, 0.5 mM ethylenediaminetetraacetic acid (EDTA), 25 units of papain per mL of solution, with CO_2_ bubbling, at 37°C for 1 h. The solution was removed and the hippocampi were incubated in 10% FBS-DMEM, 73 µM albumin for 15 minutes. The hippocampi were triturated using a 10 mL pipette in complete-neurobasal medium [Neurobasal A (Thermo Fisher), containing 2% B27 (Thermo Fisher) and 1% Glutamax-I (Thermo Fisher). Neurons were plated (12-well plate) on poly-L-lysin-hydrochloride (Sigma-Aldrich, St. Louis, Missouri, United States) coated coverslips in plating medium (500 mL MEM, 50 mL horse serum, 5 mL glutamine, 330 mg glucose. After 2 h the plating medium was replaced with 1.25 ml neurobasal-A Medium.

### Staining of BCRs

For the staining of BCRs on living cells the staining was performed on ice to avoid the internalization of BCRs. Cells (∼200,000 cells/sample) were pelleted by centrifuging at 1000 x g, resuspended in 50 µL of ice-cold complete medium (see above) containing the investigated affinity probe (see Supp. Table 1) and incubated for 10 minutes on ice. Cells were centrifuged at 400 x g at 4°C in a table top centrifuge and the excess of probe was removed. Cells were washed by resuspension in 1 ml of ice-cold Dulbecco’s Phosphate Buffered Saline (DPBS) followed by incubation on ice for 3 minutes and centrifugation at 400 x g at 4°C. The washing step was repeated 3 times to remove most of the excess of the fluorescent probes. When a secondary probe was used (see Supp. Table 1), the cells were further incubated with 50 µL ice-cold complete medium containing the secondary reagent and incubated for another 30 minutes on ice (staining controls without secondary probes were left for the same time on DPBS only). Washing was performed as described for the primary probe. After staining, cells were resuspended in 1 ml of cold DPBS and transferred to a 12 well plate (containing PLL coated coverslips). The plate was centrifuged at 500 rpm for 5 minutes at 4°C. The DPBS was carefully discarded and cells were fixed with 1 mL of 4% paraformaldehyde and 0.1% GLU in PBS for 10 minutes on ice followed by 30 minutes at room temperature. The fixative was removed and quenched by adding 1 mL of 0.1 M Glycine in DPBS and incubated at room temperature for additional 20 minutes. Finally, cells were rinsed with 1 mL DPBS and mounted on a glass slide using Mowiol (6 g glycerol, 6 ml deionized water, 12 ml 0.2 M Tris buffer pH 8.5, 2.4 g Mowiol 4–88, Merck Millipore). The staining of BCRs of fixed Ramos cells, around 200,000 cells/sample were pelleted by centrifuging at 1000 x g, resuspended in 1 mL DPBS and transferred to a single well on a 12 well plate containing PLL coated coverslips. The cells were let to sediment on the coverslips at 37°C for 1 h. DPBS was removed and cells were fixed with one of the following conditions: 10 minutes with 4% PFA, 30 minutes with 4% PFA or 30 minutes with 4% PFA and 0.1% GLU. For all fixation conditions the first 5 minutes incubation were performed on ice and the remaining fixation time at room temperature. After fixation, the quenching of reactive aldehydes was performed as described above. Cells were finally rinsed and staining was done in 1 mL DPBS containing the different probes. After staining, cells were washed 3 times with DPBS for 5 minutes at room temperature and coverslips were mounted in Mowiol.

### Imaging and Analysis of BCRs

Cells were imaged with multicolor confocal STED microscope (Abberior Instruments, Göttingen, Germany) described below. Imaging was performed using a 640 nm excitation laser and a 775 nm depletion laser. The final raw STED images were obtained after the summation of 3 successive scans. STED images of cells were analyzed using custom written MATLAB scripts (MATLAB Release 2014b, The MathWorks, Inc., Natick, Massachusetts, United States). For each cell center, the radii of two circles were manually adjusted so that the area between the circles contained all of the cell membrane. From this area, pixels were grouped by their angle to the cell center (in 360 bins of 1°) and maximum-projected to obtain the angle-dependent intensity *ŷ*_*i*_ along the membrane. The self-similarity of this function was then assessed by calculating its normalized autocorrelation

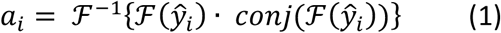

using the normalized intensity

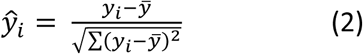

the mean value *ŷ*, the complex conjugate *conj* and the fast Fourier transform F. It gives a measure of how similar the intensity of two points are on the membrane depending on their angular distance. As the effect of different labeling homogeneities was best observed at a range of 8-12°, the autocorrelation from this area was then averaged for each cell (and translated to the perimeter in µm in the figures by approximating the cell diameters to 10 µm).

### Peroxisome size

Primary neurons from rat hippocampi were fixed with 4% PFA 30 minutes followed by 30 minutes at room temperature. The neurons were incubated in a blocking and permeabilizing solutions containing 5% bovine serum albumin (BSA) and 0.1% Triton X-100 for 20 minutes at room temperature. The rabbit polyclonal anti Pmp70 antibody (Abcam, Cat No: ab85550) was added on the cells in a 1:300 dilution in PBS containing 2.5% BSA 0.05% for 1 h at room temperature. The cells were washed 3 time for 10 minutes each in PBS and incubated with either secondary goat anti rabbit conjugated to AbberiorStar635P (Abberior GmbH, Cat. No: 2-0012-007-2) or the 2.Nb FluoTag-X2 anti rabbit also conjugated to AbberiorStar635P (NanoTag Biotechnologies, Cat. No: N1002) and diluted to 1:200 and 1:100 respectively in 2.5% BSA, 0.05% Triton X-100 for 1 h at room temperature. The cells were washed 3 times for 10 minutes in PBS and finally mounted in Mowiol. The peroxisomes on neurons were imaged with the STED setup described above using a 640 nm excitation laser and 775 nm depletion laser.

For determining the peroxisome diameter, the images were filtered using a bandpass filter, in MATLAB, to remove background noise, and peroxisome regions of interest were identified using an empiric threshold. The smallest ellipse diameter that fitted each peroxisome region of interest was then obtained by using the self-written MATLAB routine.

### Autocorrelation on Microtubule stainings

COS-7 cells were fixed with -20°C pre-cooled methanol for 20 minutes at -20°C. Methanol was removed and cells were blocked with 3% BSA for 20 minutes at room temperature. The cells were incubated with primary mouse monoclonal antibody anti-tubulin (SySy, Cat No: 302 211) directly coupled to Atto647N fluorophore and diluted at 1:25 in 1.5 % BSA for 1 h at room temperature. The cells were washed 3 times, 5 minutes each with PBS. Cells were then incubated with either secondary nanobody FluoTag-X2 anti mouse conjugated to AbberiorStar580 (NanoTag Biotechnologies, Cat No: N1202) or secondary full antibody anti mouse coupled to AbberiorStar580 (Abberior, Cat. No: 2-0002-005-1) diluted at 1:100 in 1.5% BSA for 1 h at room temperature. Finally, cells were washed as described above and mounted in Mowiol. Images of microtubules were taken using the Abberior Expert line STED system. A 640 nm excitation laser and 775 nm depletion laser were used for imaging the 1.Ab (AbberiorStar635P) while a 561 nm excitation laser and 775 nm depletion laser were used for imaging the fluorophore on the secondary probes (AbberiorStar580). The correlation of the STED signal provided by the secondary probe to the primary probe was analyzed as follows. Lines were drawn following the stained microtubules using a self-written routine in MATLAB. The Pearson‘s correlation between the directly labeled 1.Ab and the secondary probes were measured at the drawn lines. The autocorrelation of the signal from the 1.Ab was used as control.

### Pre-mixing experiment Immunostaining

COS-7 cells were fixed in -20°C pre-cooled methanol for 20 minutes at -20°C. The cells were blocked by addition of 3% BSA in PBS for 30 minutes at room temperature. In the meantime, the 1.Abs were pre-mixed for 30 minutes with two molar excess of fluorescently-labeled 2.Nbs in PBS containing 1.5% of BSA (see Supp. Table 2). The pre-mixed complexes were then incubated on the fixed cells sequentially. In between each round of pre-mixed complex, the cells were washed 3 times for 5 minute each with PBS and post fixed with 4% PFA for 10 minutes. The excess of fixative was quenched with 0.1 M glycine in PBS for 10 minutes. The cells were mounted in Mowiol and imaged using a multicolor laser scanning confocal microscope (the STED system described before).

### Pre-mixing experiment Western Blot

A confluent plate of COS-7 cells was briefly washed with ice-cold PBS before lysing the cells on the plate sitting on ice with pre-chilled Lysis buffer (50 mM Tris/HCl, pH 7.5, 150 mM NaCl, 2 mM EDTA, 0.5% IgePAL, 0.5 % Sodium deoxycholate 33and freshly added DNAse, 1 mM PMSF and protease inhibitor cocktail (Roche). Cells were scrapped and passed through a syringe with needle gauge 26 several times avoiding foam. After max. speed centrifugation at 4°C in a table-top centrifuge for 15 minutes. Supernatant was taken and mixed with 2x loading dye (50 mM Tris-HCl, 4% sodium dodecyl sulfate (SDS), 0.01% Serva Blue G, 12% glycerol, pH 6.8, 50 mM DTT) and heated at 95°C for 10 minutes. Boiled samples were then loaded in 10% SDS-PAGE. Proteins in the gel were then transferred to a nitrocellulose membrane in wet trans-blot cell (Biorad). The membranes were blocked in blocking buffer (5% Nonfat Dried Milk in PBS + 0.1% Tween-20) for 1 hour at room temperature. 1.Abs were pre-mixed with the corresponding fluorescent 2.Nb for 10 minutes and then added together on the blocked nitrocellulose membranes for 60 minutes at room temperature. Membranes were washed 5 times with large volumes of PBS for 5 minutes each and read with a LiCor Sytem Odyssey Clx.

### DNA coupling of nanobodies

Secondary nanobodies (obtained from NanoTag Biotechnologies GmbH) were coupled to docking oligonucleotide strands (Biomers GmbH, Ulm, Germany) functionalized with an azide group at the 5’-end and an Atto488 fluorophore at the 3’-end following the protocol described by Sograte-Idrissi et al^26^. In brief, the nanobody containing an extra C-terminal cysteine was reduced with 5 mM TCEP (Sigma-Aldrich, Cat. No. C4706) for 2 h on ice. TCEP was removed via 10 kDa molecular weight cut-off (MWCO) Amicon spin filters (Merck, Cat. No. UFC500324) and the nanobody was coupled through maleimide conjugation chemistry to a maleimide-DBCO crosslinker (Sigma-Aldrich, Cat. No. 760668). After removal of excess crosslinker through 10 kDa MWCO Amicon spin filters, the nanobody was coupled to the docking oligo containing an azide group at its 5‘-end (Biomers) through a strain promoted azide-alkyne cycloaddition reaction. To avoid background signal, the excess of docking oligo was removed by a size exclusion chromatography column (Superdex® Increase 75, GE Healthcare) on an Äkta pure 25 system (GE Healthcare). The docking strand sequences were obtained from Agasti el al.^24^ and can be found in Supp. Table 2.

### DNA-coupling of antibody

Donkey anti-mouse secondary antibody (Jackson Immunosearch, Cat. No. 715-005-151) was labeled with a DNA strand via a DBCO-sulfo-NHS ester linker according to the protocol as previously described by Schnitzbauer *et al*. (please cite nature protocols). Briefly, primary amines of the antibody were reacted with a DBCO-sulfo-NHS ester cross-linker (Jena Bioscience, Cat. No. CLK-A124-10) for two hours at 4°C. Unreacted cross linker was then removed using a Zeba desalting column (40 kDa MWCO, Thermo Fisher Scientific, Cat. No. 87766). The antibody-DBCO conjugate was then attached to a DNA strand functionalized with an azide group at the 5’-end via copper-free click chemistry. Excess DNA-strands were removed using 100 kDa MWCO Amicon spin filters (Merck Millipore, Cat. No. UFC510096). Docking strand sequences were obtained from Agasti el al.^24^ and can be found in Supp. Table 2.

### Stainings for DNA-PAINT

Cells for DNA PAINT imaging were plated on an 8-well chamber coverglass II (Sarstedt, Cat No: 94.6190.802 or ibidi, Cat. No. 80827 ibidi, Cat. No. 80827) and grown overnight.

The next day, cells were fixed. Cos7 cells were fixed with pre-cooled methanol for 20 minutes at -20°C. The cells were then blocked with 3% (w/v) BSA for 20 minutes at room temperature and incubated with a primary mouse monoclonal anti tubulin antibody directly labeled with Atto647N (SySy, Cat No: 302 211) and diluted 1:25 in 1.5% BSA for 1 h at room temperature. Unbound 1.Ab was removed by washing the cells 3 times with PBS for 5 minutes each. They were then incubated with the 2.Nb or 2.Ab coupled to DNA-PAINT docking sequences. The cells were washed 3 times for 5 minutes with PBS.

Rat primary hippocampal neuron, were fixed by adding 4% PFA for 30 minutes on ice and 4% PFA for 30 additional minutes at room temperature. The neurons were blocked and permeabilized with 3% (w/v) BSA + 0.1% (v/v) Triton X-100 for 20 minutes at room temperature. The mouse monoclonal anti Bassoon (Enzo, Cat No: ADI-VAM-PS003-F) and the mouse monoclonal anti Homer (SySy, Cat No: 1600111) were pre-mixed in a 1:5 molar ratio with 2.Nb anti mouse coupled to P1 (5‘-TTATACATCTATTTT-Atto488-3’) and P5 (5‘-TTTCAATGTATTTTT-Atto488-3`) respectively. The pre-mixed anti Homer 1.Ab and its 2.Nb were added on the cells for 1 h with slow orbital shaking. The cells were then washed 3x 5 minutes each with PBS and 1x 5 minutes with PBS supplemented with 0.1 M NaCl. The 1.Ab-2Nb complex were briefly fixed by adding 4% PFA for 5 minutes. The fixative was removed and the remained quenched with 0.1 M glycine for 5 minutes. The pre-mixed anti Bassoon 1.Ab with its 2.Nb was added to the cells for 1 h at room temperature, and post-fixed and quenched as before. For drift correction purposes, cells were incubated with a 1:10 dilution of 90 nm gold particles (cytodiagnostics, Cat. No. G-90-100) for 10 minutes, rinsed 4x with PBS and stored at 4°C until imaging was performed.

### DNA-PAINT Imaging

The correspondent imager strand to the DNA-PAINT docking sites used on the nanobodies (Supp. Table 3), were equipped with a Cy3b fluorophore at their 3’-end. Imager strands were diluted in PBS supplemented with 500 mM NaCl and 1x Trolox (Sigma-Aldrich, Cat. No. 238813-1G). Imager strands were used at concentrations between 0.5 nM and 2 nM to optimize the number of binding events per time (see Supp. Table 4). The focal plane was found by searching in the 488 nm channel. Cells were then imaged in the 561 nm channel with a 100-200 ms exposure time per frame for 30.000-60.000 frames. When exchange of imager was performed, the chamber was washed 10 times with PBS supplemented with 0.5 M NaCl until no residual blinking was observed anymore. The reconstruction of the raw data and the drift correction with cross correlation and gold particles as fiducial markers was performed with Picasso Sotware^4^. Microtubule filament sizes were measured via exported regions and Gaussian fits in Origin on the localizations. Images were acquired as described below and raw data movies were reconstructed with the Picasso software suite. Drift correction and multicolor alignment was performed via redundant cross-correlation and 90 nm gold particles as fiducial markers.

### Cochlear staining

Mice C75Bl6/J of 3 weeks of age were euthanized by decapitation. Cochleae were harvested and fixed in 4% PFA for 45 minutes at room temperature. Afterwards, they were processed following the cochlea-adapted version of the iDISCO+ protocol (Keppeler and Duque-Afonso et al., in preparation). Briefly, they were decalcified in 10% EDTA in PBS, pH 8, for 2 days and treated with 25% N,N,N’,N’-Tetrakis(2-Hydroxypropyl)ethylenediamine in PBS for another 2 days, in order to remove endogenous fluorescence^44^ at room temperature under constant rotation. The samples underwent the methanol-free pretreatment of the iDISCO+ protocol^39^, followed by the regular procedure for immunostaining and clearing. The pretreatment consisted in subsequent incubations at 37°C under constant shaking of the following solutions: 0.2% Triton X-100 in PBS (2×1h), 0.2% Triton X-100/20% DMSO in PBS (1 day), 0.1% Triton X-100/20% DMSO/0.1% Tween-20/0.1% Deoxycholate/0.1% IGEPAL CA-630 in PBS (1 day), Triton X-100 in PBS (2×1h). The immunostaining continued at 37°C, under constant shaking, with the incubation of the tissue in a Permeabilization solution (0.16%TritonX-100/20%DMSO/2.3% Glycine (0.3M) in PBS, 2 days) and in a Blocking Solution (0.16% TritonX-100/10%DMSO/3%BSA in PBS, 2 days). The 1.Ab (Guinea Pig antiserum anti-parvalbumin-α, 195 004, Synaptic System) was pre-mixed with the 2.Nb (Nanobody anti-guinea pig Alexa 546) using a molar ratio of 1:3 or 45 min, under constant rotation, at room temperature. The PTwH buffer contained 0.2% Tween-20/0.001% Heparin in PBS. The primary antibody was diluted in a solution containing 5%DMSO/1.5%BSA in PTwH with a concentration of 1:300. The 2.Ab (Goat-Anti Guinea pig 568, Invitrogen, A11075, 1:500) and the 1.Ab pre-mixed with the 2.Nb were diluted in a solution containing only 1.5%BSA in PTwH. The sample were incubated in 4 different ways (37°C, under shaking): 1) 6 days and 2) 14 days in the solution containing the 1.Ab premixed with the 2.Nb, 3) 3 days and 4) 7 days with the 1.Ab followed by a washing step of 1 day in PTwH at room temperature and the incubation of the 2.Ab for 3 and 7 days respectively. Before the clearing procedure, the samples were washed in PTwH for 1 day at room temperature. Finally, samples were dehydrated in an increasingmethanol dilution series (20, 40, 60, 80, 100 and 100% Methanol in ddH2O, one hour each), incubated in 66% Dicloromethane/33% Methanol for 3 hours plus two consecutive incubation in 100% DCM for 15 minutes each for lipid extraction, and immerse in Dibenzylether, as a refractive index matching solution.

### Cochlear probe penetration quantification

The original stack was resampled by a factor of 2.15×2.15×2 and converted to 8-bits in FIJI. Then, the ganglion was coarsely segmented manually with TrakEM2^45^ and imported to 3DSlicer^46,47^. There, a median filter with a kernel of 10×10×1 pixel was applied and the resulting image was threshold segmented, converted to a 3D closed surface or mesh and stored as a .stl file, as it is the input format needed for the following step. Centerlines of the ganglion were then calculated using the vmtkcenterline function of the open source software VMTK (the Vascular Modelling Toolkit, Orobix Srl) and then imported to MATLAB for further analysis. For every sample, the mesh, centerline and raw stack were imported to MATLAB. The centerline was fitted using spline interpolation and 100 position equally spaced were retrieved. In each of these positions, 14 radius of 200 µm were positioned, 6 orthogonal to the rest. The chosen orientation was parallel to the apical-basal axis formed by the most apical and most basal coordinate of the centerline. Those radii that were inside of the mesh, checked by the function inpolyhedron^48^, or outside of the original image space, were removed. Radii were mapped in the image space and the pixel values in their coordinates were used to obtain the line profiles. The minimum of each profiles was subtracted for each to have a comparable baseline.

### Microscopy Setups

Fluorescent imaging of Supp Fig.1 was done with Nikon inverted epifluorescence microscope. The microscope was equipped with an HBO 100-W lamp and an IXON X3897 Andor Camera. For all samples, a 60X Plan apochromat oil immersion objective (NA 1.4) was used (from Nikon). The filter sets and time course (if applicable) used for imaging are shown in Table 3. Images were obtained using the image acquisition software NiS-Elements AR (Nikon).

STED microscopy images were obtained using STED Expert line microscope (Abberior Instruments, Göttingen, Germany) composed of a IX83 inverted microscope (Olympus, Hamburg, Germany) with a UPLSAPO 100x 1.4 NA oil immersion objective (Olympus). Confocal images were obtained from the same setup without using the STED depletion laser.

DNA-PAINT imaging was carried out on an inverted Nikon Eclipse Ti microscope (Nikon Instruments) with the Perfect Focus System, applying an objective-type TIRF configuration with an oil-immersion objective (Apo SR TIRF 100×, NA 1.49, Oil). Two lasers were used for excitation: 561nm (200 mW, Coherent Sapphire) or 488 nm (200 mW, Toptica iBeam smart). The laser beam was passed through a cleanup filter (ZET488/10x or ZET561/10x, Chroma Technology) and coupled into the microscope objective using a beam splitter (ZT488rdc or ZT561rdc, Chroma Technology). Fluorescence light was spectrally filtered with two emission filters (ET525/50m and ET500lp for 488 nm excitation and ET600/50 and ET575lp for 561 nm excitation, Chroma Technology) and imaged on a sCMOS camera (Andor Zyla 4.2) without further magnification, resulting in an effective pixel size of 130 nm after 2×2 binning. Camera Readout Sensitivity was set to 16-bit, Readout Bandwidth to 540 MHz.

Light sheet images of the cochleae were done using a lightsheet microscope (LaVision Biotec Ultramicroscope II). The laser power was constant for all the samples except for the sample incubated with 1.Ab-2.Ab for 14 days, which was 6.75 times lower (13.5% vs. 2%). The stacks were acquired with a total zoom of 8x (2x MVPLAPO Objective and 4x Optic Zoom microscope body), a stepsize of 3 µm, with a lightsheet of 30% width and a thickness of 5 µm (NA: 0.148, unidirectional illumination and 11-12 steps of dynamic horizontal focus. The images were imported to FIJI^49^ for calculating the maximum intensity projection image and to generate the RGB tif files with a mpl-magma look-up-table.

## Acknowledgments

We thank Niklas Engels for providing Ramos cells. We thank Riccardo Testolin for the help in designing the hybrid plot graphs. We thank Eugenio F. Fornasiero, Sebastian Jaehne and Sven Truckenbrodt for reading and commenting the manuscript. This work was supported by the Deutsche Forschungsgemeinschaft (DFG) through Cluster of Excellence Nanoscale Microscopy and Molecular Physiology of the Brain (CNMPB) to F.O and the Cluster of Excellence Multiscale Bioimaging: from Molecular Machines to Networks of Excitable Cells (MBExC). T.S. and S.S. acknowledge support from the DFG through the Graduate School of Quantitative Biosciences Munich (QBM). This work was further supported by the DFG through the Emmy Noether Program (DFG JU 2957/1-1), the SFB 1032 (Nanoagents for spatiotemporal control of molecular and cellular reactions, Project A11), the ERC through an ERC Starting Grant (MolMap, grant agreement no. 680241), the Max Planck Society, the Max Planck Foundation, and the Center for Nanoscience (CeNS) to R.J.

## Author contributions

S.SI. designed and performed experiments, analyzed data, and wrote the manuscript, T.S. and S.S. designed and performed experiments, analyzed data and revised the manuscript. C.DA designed and performed experiments, analyzed data and revised the manuscript. M.A. created image analysis macros and revised the manuscript. T.M designed experiment and revised the manuscript. R.J. designed experiment and revised the manuscript. S.O.R. designed experiments, analyzed data and revised the manuscript F.O. designed and conceived the project, designed and performed experiments and wrote the manuscript.

