## Supplementary material for "Circumvention of common labeling artifacts using secondary nanobodies": Supp. Fig.

### Supplementary Table 1 | Antibody used

| Probe name | Company | Catalogue number | Dilution used |
|---|---|---|---|
| affibody® anti-IgM coupled to the Star635P | Abcam, Cambridge, UK | ab36088 | 1:25 |
| Anti-IgM polyFab' coupled to the Star635P | Jackson ImmunoResearch, Cambridgeshire, UK | cleaved with Papain from 109-006-129 | 1:50 |
| Mouse monoclonal anti IgM | Abcam, Cambridge, UK | ab193159 | 1:200 |
| Secondary donkey anti rabbit-Star635P | Abberior, Goettingen, Germany | 2-0012-007-2 | 1:200 |
| Secondary nanobody anti rabbit-Star635P; FluoTag-X2 anti Rabbit | NanoTag Biotechnology, Goettingen, Germany | N1002-Ab635P | 1:50 |
| Monoclonal mouse Anti-GM130 | BD bioscience | 610822 | 1:62,5 |
| Monoclonal mouse anti NPC | Abcam, Cambridge, UK | ab24609 | 1:200 |
| Mouse monoclonal anti alpha Tubulin | Synaptic Systems, Goettingen, Germany | 302211 | 1:500 |
| Secondary nanobody anti mouse FluoTag-X2 anti Mouse kLC | NanoTag Biotechnology, Goettingen, Germany | N1202 | 1:100 |
| Secondary donkey anti-mouse antibody | Jackson ImmunoResearch, Cambridgeshire, UK | 715-005-151 | 1:100 |
| Monoclonal mouse anti Beta actin | Sigma-Aldrich, Missouri, USA | A1978 | 1:100 |
| Polyclonal rabbit anti Lamin B | Sigma-Aldrich, Missouri, USA | HPA050524) | 1:100 |
| FluoTag-X2 anti Mouse kLC LiCor800CW | NanoTag Biotechnology, Goettingen, Germany | N1202-Li800 | 1:500 |
| FluoTag-X2 anti Mouse kLC LiCor680RD | NanoTag Biotechnology, Goettingen, Germany | N1202-Li680 | 1:500 |
| FluoTag-X2 anti Rabbit LiCor800CW | NanoTag Biotechnology, Goettingen, Germany | N1202-Li800 | 1:500 |

### Supplementary Table 2 | Handle sequences

| Handle Name | Sequence | 5'-mod | 3'-mod | Company |
|---|---|---|---|---|
| P1 | TTATACATCTATTTT | Azide | Atto488 | Biomers.net |
| P3 | TTTCTTCATTATTTT | Azide | Atto488 | Biomers.net |
| P5 | TTTCAATGTATTTTT | Azide | Atto488 | Biomers.net |

### Supplementary Table 3 | Imager sequences

| Imager name | Sequence | 5'-mod | 3'-mod | Company |
|---|---|---|---|---|
| P1* | CTAGATGTAT | None | Cy3b | Eurofins Genomics |
| P3* | GTAATGAAGA | None | Cy3b | Eurofins Genomics |
| P5* | CATACATTGA | None | Cy3b | Eurofins Genomics |

**Supplementary Table 4| Imaging parameters**

| Dataset | Parameters | Power @561 nm |
|---|---|---|
| Figure 2A-C: DNA-PAINT Microtubule secondary nanobody | 200ms, 2D, 60k Frames, 2nM. P1* | 1 $kW/cm^2$ |
| Figure 2D-F: DNA-PAINT Microtubule secondary antibody | 200ms, 2D, 60k Frames, 2nM. P1* | 1$kW/cm^2$ |
| Figure 3C-E: Bassoon | 150ms, 3D, 30k Frames, 3nM, P5* | 1 $kW/cm^2$ |
| Figure 3C-E: Homer | 150ms, 3D, 30k Frames, 6 nM, P3* | 1 $kW/cm^2$ |

**Supplementary Table 5|** Statistics on BCR autocorrelation Analysis. One-way Anova with Tukey Multiple Comparison Test. ns= non-significant, *= $p \leq 0.05$, **= $p \leq 0.01$, ***= $p \leq 0.001$, ****= $p \leq 0.0001$

| | polyFab' live | Affibody live | 1.Ab+2.Ab live | 1.Ab+2Nb live |
|---|---|---|---|---|
| **polyFab' live** | | ns | **** | ** |
| **Affibody live** | | | **** | ns |
| **1.Ab+2.Ab live** | | | | *** |
| | **1.Ab+2.Ab live** | **1.Ab+2.Ab 10 min 4% PFA** | **1.Ab+2.Ab 30 min 4% PFA** | **1.Ab+2.Ab 30 min 4% PFA+ 0.1% GLU** |
| **1.Ab+2.Ab live** | | ns | *** | **** |
| **1.Ab+2.Ab 10 min 4% PFA** | | | * | ns |
| **1.Ab+2.Ab 30 min 4% PFA** | | | | ns |
| | **1.Ab+2Nb live** | **1.Ab+2Nb 10 min 4% PFA** | **1.Ab+2Nb 30 min 4% PFA** | **1.Ab+2Nb 30 min 4% PFA+ 0.1% GLU** |
| **1.Ab+2Nb live** | | ns | ns | ns |
| **1.Ab+2Nb 10 min 4% PFA** | | | ns | ns |
| **1.Ab+2Nb 30 min 4% PFA** | | | | ns |
| | **polyFab' live** | **polyFab' 10 min 4% PFA** | **polyFab' 30 min 4% PFA** | **polyFab'30 min 4% PFA+ 0.1% GLU** |
| **polyFab' live** | | ns | ns | ns |
| **polyFab' 10 min 4% PFA** | | | ns | ns |
| **polyFab'30 min 4% PFA** | | | | ns |

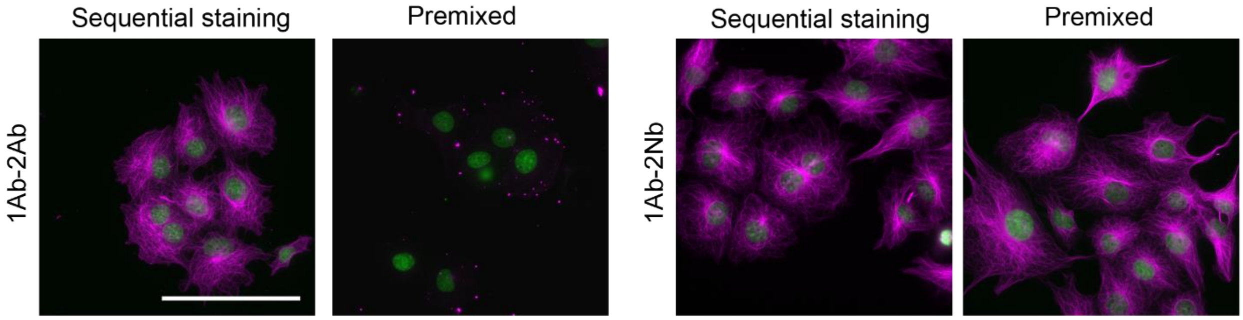


**Supplementary Figure 1**: Pre-mixing antibodies in a centrifuge tube prior incubation on the cell. Immunostaining is commonly done by sequential incubation of the primary probe and the secondary probe. Pre-mixing the two probe in a centrifuge tube prior incubation leads to no staining for 1.Ab-2.Ab while staining is maintained for 1.Ab-2.Nb. Hoechst staining (nucleus) in green, microtubule staining in magenta. Scale bar= 50 µm.

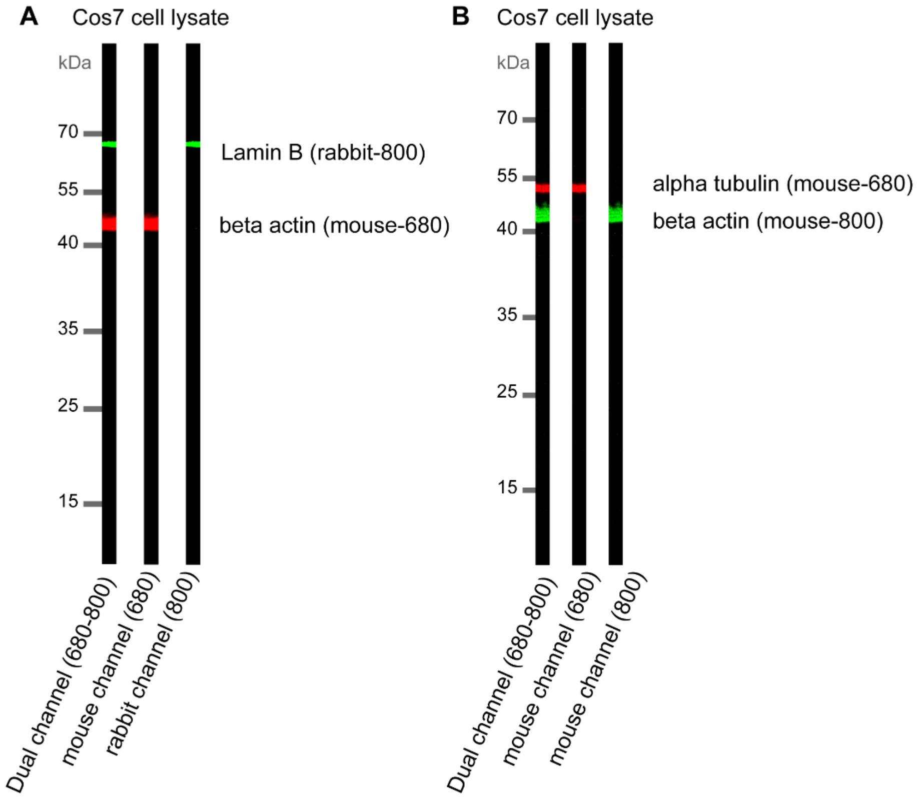


**Supplementary Figure 2**: Pre-mixing 1.Ab-2.Nb for Western Blot. COS-7 cell lysate blotted on nitrocellulose membrane. **A)** Pre-mixing allows shorter protocol by one single step staining. The membrane was stained with 1.Ab beta actin pre-mixed with 2.Nb anti Mouse coupled to IRDye680RD and 1.Ab anti Lamin B pre-mixed with 2.Nb anti

Rabbit-IRDye800CW **B)** Pre-mixing allows use of same species antibodies in the same westen blot. The membrane was stained with 1.Ab beta actin pre-mixed with 2.Nb anti Mouse-IRDye800CW and 1.Ab anti alpha tubulin pre-mixed with 2.Nb anti Mouse-IRDye800CW

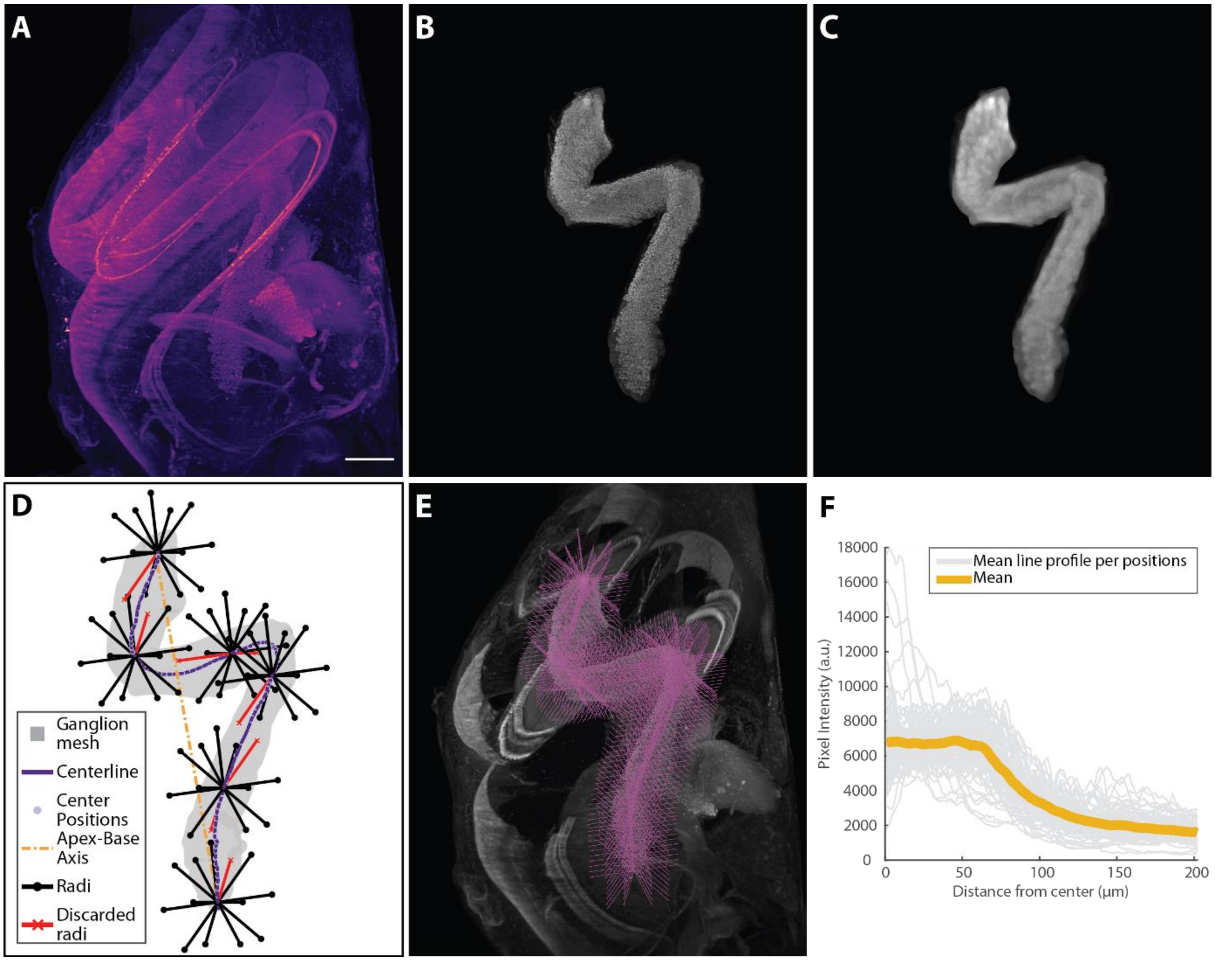


**Supplemental Figure 3.** Method to investigate the sample penetration of different labelling approaches in cochlear staining**. A)** Maximal intensity projection of a cleared cochlea stained with 1.Ab against parvalbumin-$\alpha$ premixed with 2.Nb anti-guinea pig. **B)** Coarse manual segmentation of the ganglion**. C)** Median filtered image of the ganglion (kernel: 10x10x1). **D)**. 2D projection of the mesh created from a threshold segmentation of C), its centerline, the apex-base axis, the center positions where the radii fan out and the used and discarded radii. Only 6 out of the 100 center positions and their corresponding radii used are displayed for clarity. **E)** Maximal intensity projections of a sub-stack of the slices that contains only the ganglion. In magenta, all the radii mapped back in the image space. **F)**. Mean line profile per position (n=100 positions) and mean line profile for this sample is plotted against the distance from the center position. Scalebar for A-C and E: 200 µm.

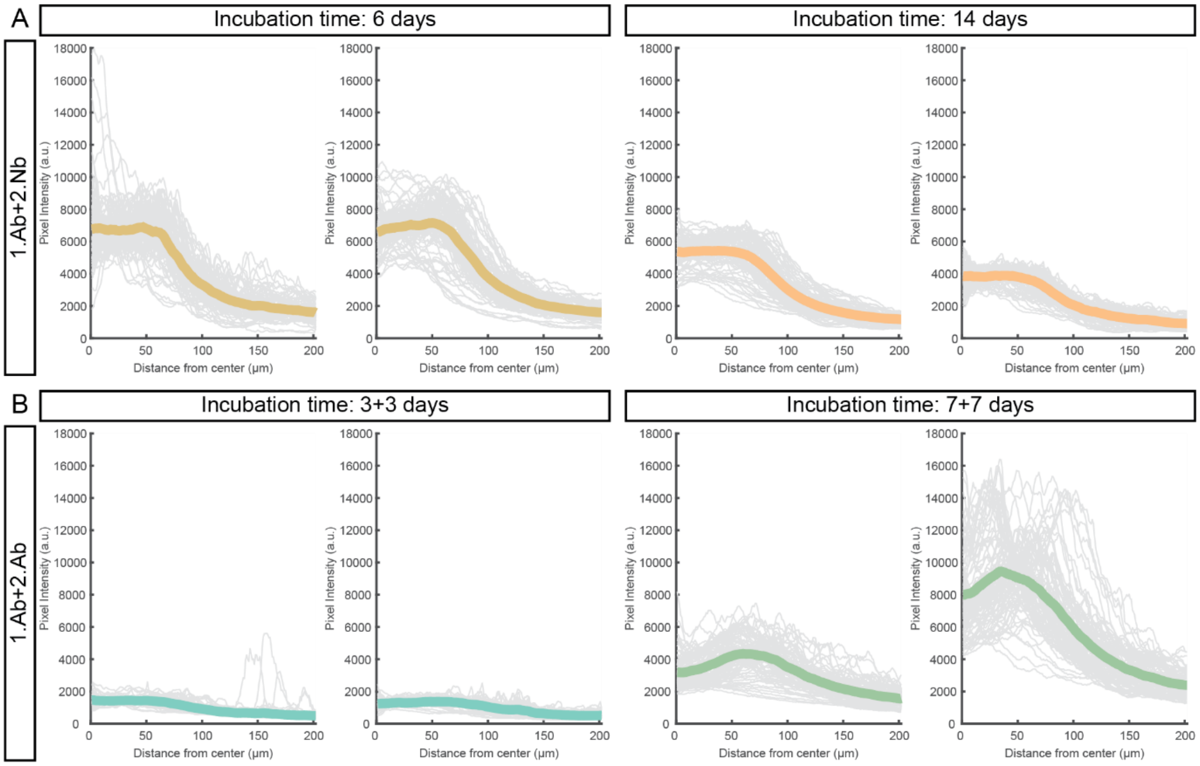


**Supplemental Figure 4. Line profile from individual cochlear samples.** Mean profile per position (n= 100 per sample, grey thin traces) and mean profile per sample (N=2 per staining method and incubation time, color thick traces) are displayed against distance from center position from **A)** Samples stained with a 1.Ab against parvalbumin-$\alpha$ premixed with 2.Nb against guinea pig, labeled with Alexa Fluor 546, and **B)** Samples stained with a 1.Ab against parvalbumin-$\alpha$ revealed by a 2.Ab against guinea pig, labeled with Alexa Fluor 568.

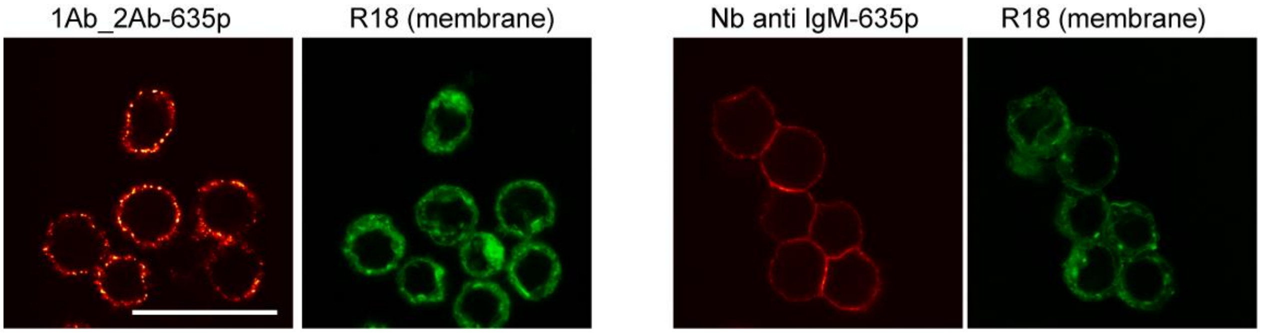


**Supplementary Figure 5**: Diffraction limited images (confocal microscope) of B cells stained with 1.Ab-2.Ab (left panel) or primary nanobody 1Nb (right panel) targeting the IgM of the BCR receptor. In green a membrane staining is performed (R18) to show the integrity of the membrane. z

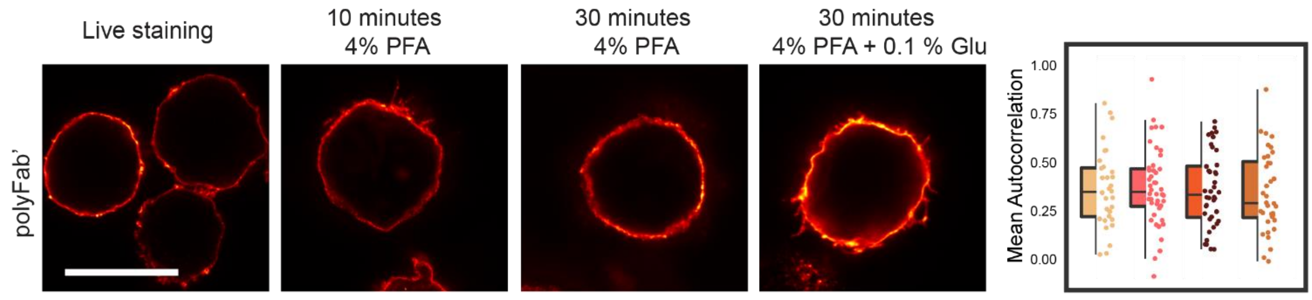


**Supplementary Figure 6:** B cells fixed in different conditions and subsequently stained with polyFab'. STED images and autocorrelation analysis as explained in Fig.4.

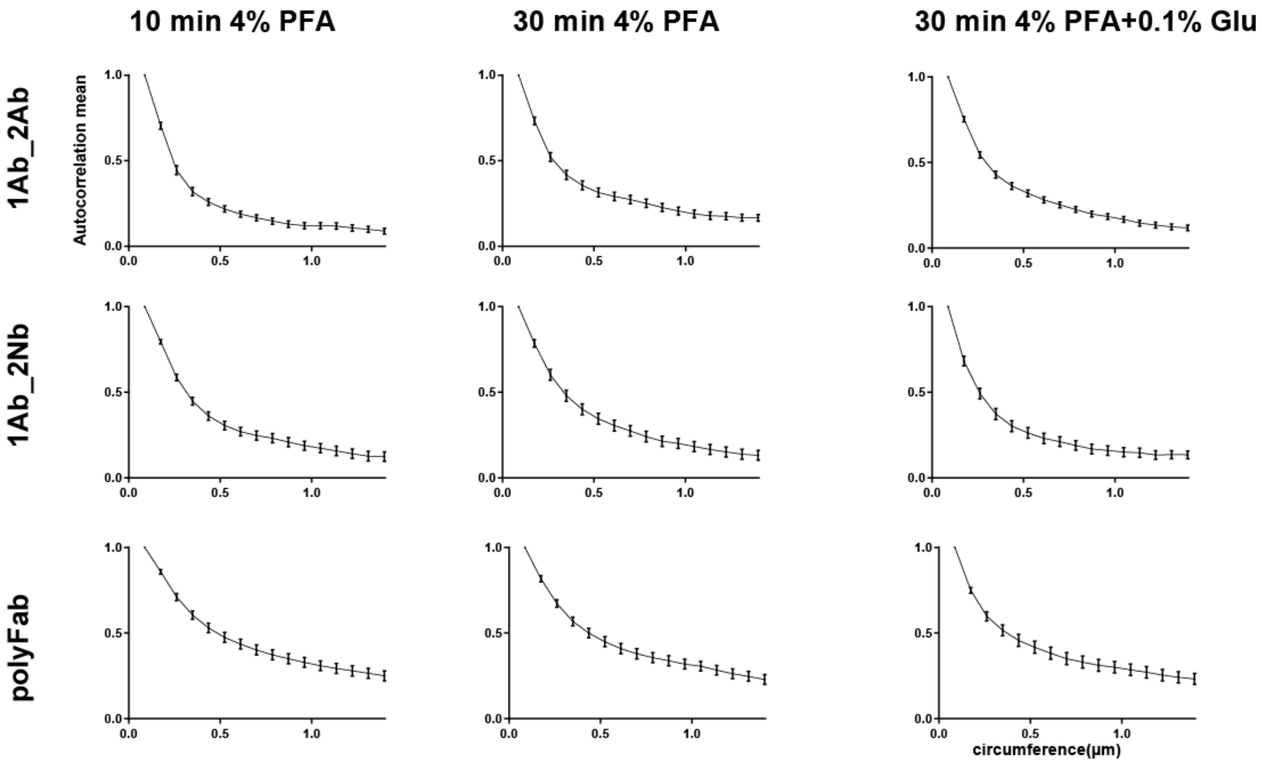


**Supplementary Figure 7**: Autocorrelation curve of B cells fixed prior staining with different fixation conditions. Selected images and analyses are in Figure1 (d-e)
